# Root volatiles in plant-plant interactions I: Characterization of root sesquiterpene emissions from *Centaurea stoebe* and their effects on other plants

**DOI:** 10.1101/441717

**Authors:** Valentin Gfeller, Meret Huber, Christiane Förster, Wei Huang, Tobias G. Köllner, Matthias Erb

## Abstract

Volatile organic compounds (VOCs) emitted by plant leaves can influence the physiology of neighboring plants. In contrast to interactions above ground, little is known about the role of VOCs in belowground plant-plant interactions. Here, we characterize constitutive root volatile emissions of the spotted knapweed (*Centaurea stoebe*) and explore the impact of these volatiles on the germination and growth of different sympatric plant species. We show that *C. stoebe* roots emit high amounts of sesquiterpenes, with estimated release rates of (*E*)-β-caryophyllene above 3 μg g^−1^ dw h^−1^. Sesquiterpene emissions show little variation between different *C. stoebe* populations, but vary substantially between different *Centaurea* species. Through root transcriptome sequencing, we identify six root-expressed sesquiterpene synthases (TPSs). Two root-specific TPSs, CsTPS4 and CsTPS5, are sufficient to produce the full blend of emitted root sesquiterpenes. Volatile exposure experiments demonstrate that *C. stoebe* root volatiles have neutral to positive effects on the germination and growth of different sympatric neighbors. Thus, constitutive root sesquiterpenes produced by two *C. stoebe* TPSs are associated with facilitation of sympatric neighboring plants. The release of root VOCs may thus influence *C. stoebe* abundance and plant community structure in nature.

## Introduction

Plants influence their environment to maximize their fitness. One strategy by which plants can manipulate their environment is to produce and release chemicals such as volatile organic compounds (VOCs) (Pichersky & Gang 2000). VOCs can for instance protect plants against biotic and abiotic stress (Gouinguené & Turlings 2002; Loreto & Schnitzler 2010; Pichersky & Gershenzon 2002; Peñuelas et al. 2014). VOCs can also influence defense and growth of neighboring plants (Karban, Yang & Edwards 2014; Kegge et al. 2015; Ninkovic 2003; Pierik et al. 2003). Although the benefits of VOC-mediated plant-plant interactions for the emitter are subject to debate (Heil 2014; Morrell & Kessler 2017), VOC-mediated plant-plant interactions are increasingly recognized to influence plant ecology in natural and agricultural systems (Ninkovic, Markovic & Dahlin 2016). While most work on plant VOCs has focused on the phyllosphere, an increasing number of studies demonstrate that plant VOCs also have important roles in the rhizosphere. Root VOCs can for instance influence the behavior of herbivorous insects (Robert et al. 2012) and nematodes (Rasmann et al. 2005) and affect soil bacterial and fungal communities (Kleinheinz et al. 1999; Wenke, Kai & Piechulla 2010). In petri dish experiments, root VOCs have also been shown to negatively affect seed germination and seedling growth (Ens et al. 2009; Jassbi, Zamanizadehnajari & Baldwin 2010). Whether root VOCs mediate plant-plant interactions under more realistic conditions remains to be determined (Delory et al. 2016).

With more than 30,000 different structures, terpenoids are the most diverse class of secondary metabolites in the plant kingdom (Hartmann 2007) and are an integral part of plant VOC blends (Gershenzon & Dudareva 2007). Most volatile terpenoids are hemiterpenes (C_5_), monoterpenes (C_10_) and sesquiterpenes (C_15_) (Nagegowda 2010). Volatile terpenes have various ecological effects and function in plant-plant, plant-insect and plant-microbe interactions (Cheng et al. 2007). Terpenoids are derived from two common C_5_ precursor molecules, isopentenyl diphosphate (IPP) and its allylic isomer dimethylallyl diphosphate (DMAPP). In higher plants, IPP and DMAPP are formed through two different pathways, the mevalonic acid (MVA) and the methylerythritol phosphate (MEP) pathway. IPP and DMAPP are then further converted into geranyl diphosphate (GPP) and farnesyl diphosphate (FPP) as precursors for mono- and sesquiterpenes, respectively. The reaction for the final conversion to mono- and sesquiterpenes is catalyzed by terpene synthases (TPSs), which require a divalent cation to mediate the terpene formation (Cheng et al. 2007; Nagegowda 2010). As key enzymes for the production of terpenes, TPSs have been characterized in plants (Degenhardt, Köllner & Gershenzon 2009; Jia et al. 2018), insects (Beran et al. 2016), fungi (Quin, Flynn & Schmidt-Dannert 2014), bacteria (Yamada et al. 2015), and amoebae (Chen et al. 2016). In plants it is known that TPS expression can be regulated in a tissue specific manner. Furthermore, TPSs often catalyze the formation of multiple products, which contributes to the substantial structural diversity of terpenoids (Tholl 2006).

In this study we characterize root VOCs emitted by the spotted knapweed (*Centaurea stoebe*). The tetraploid cytotype of *C. stoebe* is invasive in northern America (Treier et al. 2009), whereas the diploid cytotype is classified as threatened (vulnerable) in Switzerland according to the International Union for Conservation of Nature (IUCN). A previous study found that *C. stoebe* root chemicals affect the physiology of *Taraxacum officinale* agg. roots and their suitability for root feeding *Melolontha melolontha* larvae (Huang et al. 2018). As no direct root contact was needed to trigger these effects, we hypothesized that *C. stoebe* may affect neighboring plants through the release of root VOCs. In this study, we analyze the volatile blend of *C. stoebe* roots and identify sesquiterpenes as dominant root VOCs. Through root transcriptome sequencing and heterologous expression, we identify TPSs that are associated with this phenotype. Furthermore, we assess the impact of *C. stoebe* roots on the germination and growth of different sympatric plant species. The results of this study also provide a mechanistic basis to determine the impact of *C. stoebe* root sesquiterpenes on *T. officinale* and its interaction with *M. melolontha* larvae (companion paper Huang et al., under review). This work thus sheds light on the genetic basis and ecological consequences of VOC-mediated plant-plant interactions below ground.

## Methods and Materials

### Study system

*Centaurea stoebe* L. (diploid) plants were grown from seeds purchased from UFA-SAMEN (Winterthur, Switzerland), unless specified otherwise. Seeds of *Anthemis tinctoria* L., *Centaurea scabiosa* L., *Centaurea jacea* L., *Cichorium intybus* L., *Daucus carota* L., *Dianthus carthusianorum* L., *Echium vulgare* L., *Festuca valesiaca* Gaudin, *Ranunculus bulbosus* L., *Taraxacum officinale* agg were obtained from the same vendor. *Medicago sativa* L. was obtained from Sativa Rheinau AG (Rheinau, Switzerland) and *Cardaria draba* (L.) Desv., was obtained from Templiner Kräutergarten (Templin, Germany). *Centaurea valesiaca* (DC.) Jord. seeds were collected from a natural population in Raron (VS, Switzerland) and provided by Adrian Möhl (Info Flora) and Markus Fischer (University of Bern). Two *C. stoebe* populations Hu-11 (tetraploid, Hungary) and Ro-11 (tetraploid, Romania), as well as *Koeleria macrantha* (Ledeb.) Schult. (MT, USA) were provided by Yan Sun and Heinz Müller-Schärer (University of Fribourg). Detailed information on these *C. stoebe* populations can be found in Mráz et al.(2012). Plant growth conditions are described in the corresponding experimental sections below.

### Characterization of C. stoebe root volatiles

To determine root volatile release by *C. stoebe*, plants were grown individually in sand under controlled conditions in a growth chamber (day length: 16 h; temperature: 20-22 °C; humidity: 65%) for seven weeks. The root system of each plant was then washed, separated from the shoot with a scalpel and dried with a paper towel (n = 8). Subsequently the roots were weighted and the cut at the root-shoot junction was sealed with Teflon tape before analysis to avoid contamination of the headspace with wound-released VOCs. The roots where then carefully inserted into 20 mL screw top glass vials (Gerstel, Sursee, Switzerland) and closed with airtight screw caps (septum Silicone/PTFE; Gerstel, Sursee, Switzerland). The vials were incubated for 1 min at 20 °C. Volatiles were then collected by exposing a SPME fiber (coated with 100 μm polydimethylsiloxane; Supelco, Bellefonte, PA, USA) to the headspace for 1.8 s. Volatiles were thermally desorbed (220 °C for 1 min) in the inlet of an Agilent 7820A series GC coupled to an Agilent 5977E MSD (source 230 °C, quadrupole 150 °C, ionization potential 70 eV, scan range 30–550; Palo Alto, CA, USA). After each run, the SPME fiber was baked out for 2 min at 220 °C. VOCs were separated on a capillary GC-MS column (HP5-MS, 30m, 250μm ID, 2.5μm film; Agilent Technologies, Palo Alto, CA, USA) with He as carrier gas at a flow rate of 1 mL/min. Initial column temperature was set to 60 °C for 1 min followed by three temperature gradients: (i) 7 °C/min to 150 °C, (ii) 3 °C/min to 165 °C and (iii) 30 °C/min to 250 °C and hold at this temperature for 3 min. VOCs were tentatively identified by comparing mass spectra to library entries of the National Institute of Standards and Technology (NIST 14). (*E*)-β-caryophyllene was identified by comparing mass spectrum and retention time to a synthetic standard (≥ 98.5 %, Sigma-Aldrich, Buchs SG, Switzerland). The first eluting petasitene was cross-validated by comparing mass spectra and retention times with a petasitene peak detected in a *Petasites hybridus* (L.) P. Gaertn. & al. root extract (Saritas, von Reuss & König 2002). The other petasitene-like sesquiterpenes were tentatively identified by comparing mass spectra to petasitene from *P. hybridus*.

### Quantification of terpene emissions

To quantify the emission of (*E*)-β-caryophyllene from *C. stoebe* roots, we first constructed volatile dispensers with known (*E*)-β-caryophyllene release rates. The dispensers were constructed by adding 5 μL pure (*E*)-β-caryophyllene (> 98.5%, GC, Sigma-Aldrich, Buchs SG, Switzerland) into a 0.1 mL micro-insert (15 mm top; VWR, Dietikon, Switzerland). Teflon tape was wrapped around a 1 μL capillary (Drummond, Millan SA, Plan-Les-Ouates, Switzerland), which was then plugged into the insert and sealed with more Teflon tape. The dispenser was stored for one day at room temperature before use to establish constant release rates. The (*E*)-β-caryophyllene emission rate of the dispenser was quantified as previously described (D’Alessandro & Turlings 2005). In short: the dispenser was placed into a glass bottled attached to a flow through system, whereby the outflow was coupled to a Super-Q trap to collect the volatile compounds. After 4 hours of volatile collection, the analytes were eluted from the trap with dichloromethane spiked with nonyl acetate as internal standard. The eluate was analyzed by gas chromatography-mass spectrometry (GC-MS) and compared to an (*E*)-β-caryophyllene dilution series which was directly injected into the GC-MS, thus allowing to compute the (*E*)-β-caryophyllene release rate of the dispensers. For the GC-MS analysis, 1 μL of sample was injected into the inlet of the GC-MS system followed by separation and analysis as described above. To ensure an accurate (*E*)-β-caryophyllene quantification, a single calibrated dispenser was incubated in SPME vials for different incubation periods (1, 5, 7.5, 10, 12.5, 20 min). The linear relationship between (*E*)-β-caryophyllene release and MS signal (R^2^ = 0.98) was used to calculate *C. stoebe* root (*E*)-β-caryophyllene emission. To calculate the release per g dry weight (DW), we dryed the roots after analysis (80°C for 48h) and weighed them using a microbalance (n = 8).

### Hexane tissue extraction and analysis

To analyze the composition and abundance of VOCs in *C. stoebe* root and leaf extracts, plants were grown in ‘Tonsubstrat’ (Klasmann-Deilmann, Geeste, Germany) in a greenhouse (light: 14h; temperature: day 21-23 °C night 19-21 °C; humidity: 50-60 °C) for ten weeks. Tissue samples were obtained by washing the roots and leaves, drying them with paper towel and wrapping root and leaf tissue separately into aluminum foil, flash-freezing them in liquid nitrogen and storing them at −80 °C. All samples were ground with mortar and pistil under liquid nitrogen, and approximately 100 mg of frozen tissue powder per sample were put into a 1 mL glass vial. One mL of hexane with nonyl acetate as internal standard (10 ng*μl) was immediately added to the samples (n = 10 for each tissue). The samples were shaken at 200 rpm for 1 h at room temperature, followed by a centrifugation step of 20 min at 5,300 rpm. 600 μL of supernatant per sample were pipetted into new tubes and stored at −20 °C. Characterization of VOCs in the extracts was carried out on an Agilent 6890 series GC coupled to an Agilent 5973 mass selective detector (source 230 °C, quadrupole 150 °C, ionization potential 70 eV; Palo Alto, CA, USA) and a flame ionization detector operating at 300 °C. He (MS) and H_2_ (FID) were used as carrier gases. The VOC separation took place on a DB-5MS capillary column (Agilent, Santa Clara, CA, USA, 30 m × 0.25 mm × 0.25 μm). After injection of 1 μL of tissue extract, the following temperature program was run: initial temperature of 45 °C was hold for 2 min followed by two temperature ramps, (i) 6 °C/min to 180 °C and (ii) 100 °C/min to 300 °C and hold for 2 min. For volatile quantification, the peak areas of the GC-FID chromatograms were integrated. The area of each compound was taken relative to the area of the internal standard and corrected for the weight of the extracted tissue. For compound identification, root and leaf samples were also run on the GC-MS. In parallel an n-alkane standard solution was run with the same method, which enabled to calculate the linear retention indices (RI) following the procedure published by van den Dool & Kratz (1963). Tentative identification was carried out by comparing mass spectra and RI of a given peak to known compounds in plant extracts of *Aloysia sellowii* (Briq.) Moldenke and *Phoebe porosa* (Nees & Mart.) Mez., which were kindly provided by Prof. W.A. König, University of Hamburg. For compounds not found in these plant extracts, mass spectra and RI were matched to the library entries of the National Institute of Standards and Technology (NIST 14). Corresponding retentions indices (RI) can be found in the supplementary materials (Tab. S1). Daucadiene was tentatively identified by comparison to the mass spectra in the NIST library. Although the mass spectra showed high similarity, the RI was not as described for the best match to the NIST library (trans-dauca-4(11),8-diene), suggesting that the detected compound might be another daucadiene diastereoisomer.

### Terpene emission of C. stoebe populations and related species

To study if root sesquiterpene production differs between *C. stoebe* ecotypes and between congeneric plant species, plants of three *C. stoebe* populations, as well as four different species of the genus *Centaurea* were grown in sand under controlled conditions (day length: 16 h; temperature: 20-22 °C; humidity: 65%) for five weeks. Two tetraploid populations (Hu-11, Ro-11) and one diploid population (UFA) were compared (n = 5-7). As congeneric species, *C. jacea*, *C. scabiosa* and *C. valesiaca*, which grow in distinct habitats were used (Landolt et al. 2010) (n = 4-8). Roots were prepared as descried above for VOC characterization. The glass vials containing the roots were immediately stored on a cooling block at 2 °C of an autosampler system (MPS; Gerstel, Sursee, Switzerland) connected to the GC-MS system. Immediately prior to analysis, the samples were transferred to an incubator set to 30 °C, in which VOCs were subsequently collected by exposition of an SPME fiber to the headspace for 1.8 s. Next, the compounds were analyzed on the GC-MS system as mentioned above for VOC characterization.

### Transcriptome sequencing and analysis

To explore the molecular basis of *C. stoebe* sesquiterpene production, we performed root transcriptome sequencing. *C. stoebe* root tissue was harvested, washed, dried, wrapped in aluminum foil and flash frozen in liquid nitrogen and ground to a fine powder. Total RNA was isolated from root powder following the manufactures protocol of the InviTrap^®^ Spin Plant RNA Mini Kit (Stratec molecular, Berlin, Germany). A TruSeq RNA-compatible library was prepared and PolyA enrichment was performed before sequencing the transcriptome on an IlluminaHiSeq 2500 with 10 Mio reads (250 base pair, paired end). Reads were quality trimmed using Sickle with Phred quality score of >20 and a minimum read length of 60. *De novo* transcriptome assembly was performed with the pooled reads using Trinity (version 2.2.0) running at default settings. Raw reads were deposited in the NCBI Sequence Read Archive (SRA) under the BioProject accession (to be inserted at a later date). To identify putative terpene synthase genes, the root transcriptome was screened using a TBLASTN search with the (*E*)-β-caryophyllene synthase MrTPS1 from *Matricaria recutita* (Irmisch et al. 2012) as query.

### Sequence analysis and tree reconstruction

Multiple sequence alignment of the identified TPS genes from *C. stoebe* and characterized TPS genes from *M. recutita* was computed using the MUSCLE codon algorithm implemented in MEGA6 (Tamura et al. 2013). Based on the alignment, a tree was reconstructed with MEGA6 using a maximum likelihood algorithm (GTR model). Codon positions included were 1st+2nd+3rd+noncoding. All positions with <80% site coverage were eliminated. A bootstrap resampling analysis with 1000 replicates was performed to evaluate the topology of the generated tree.

### Cloning and heterologous expression of CsTPS genes

To evaluate the TPS activity of the putative CsTPS genes, cDNA was produced. Then, focal genes were cloned into an expression vector and heterologously expressed in *Escherichia coli*. Subsequently, proteins were isolated and used for enzyme activity assays. To obtain plant material for RNA extraction, *C. stoebe* plants were grown in sand under controlled conditions (day length: 16 h; temperature: 20-22 °C; humidity: 65%) for eight weeks. Roots were gently washed, dried with a paper towel, cut 2 mm below root initiation, wrapped in aluminum foil and immediately flash frozen in liquid nitrogen. Afterwards, roots were ground with mortar and pistil under constant cooling with liquid nitrogen and stored at −80 °C before further processing. RNA extraction was carried out according to the manufacturer’s protocol with an innuPrep Plant RNA Kit (Analytik Jena, Jena, Germany). For cDNA synthesis, 2 μg of RNA was treated with DNAse (Thermo scientifics, CA, USA). First-strand DNA was synthesized with oligo dT_12-18_ primers and Super Script™ III reverse transcriptase (Invitrogen, Carlsbad, CA, USA). The open reading frames (ORF) of the putative *C. stoebe* terpene synthases were amplified with the primer pairs listed in the supplementary (Tab. S2) and cloned into a pASK-IBA37plus plasmid (IBA-Lifesciences, Göttingen, Germany) by restriction digest and ligation. NEB 10-beta competent *E. coli* cells (New England Biolabs, Ipswich, MA, USA) were then transformed with these vectors. In order to obtain the cloned *CsTPS* sequences and to check the transformation events, the inserted fragments were sequenced by Sanger sequencing.

For heterologous expression, NEB 10-beta cells containing the *CsTPS* constructs were grown at 37°C to an OD_600_ of 0.8. Subsequently protein expression was induced by adding anhydrotetracycline (IBA-Lifesciences, Göttingen, Germany) to a final concentration of 200 ng*mL^−1^. Expression took place for 18 h at 18 °C. Cells were harvested by centrifugation and resuspended in assay buffer (10 mM Tris HCl, 1mM DTT and 10 % (vol/vol) glycerol (pH 7.5)). To disrupt the cells, they were treated 4 × 20 s at 60 % power with a sonicator (Bandelin Sonoplus HD 2070, Berlin, Germany). Samples were then centrifuged at 4 °C for 1 h at 14,000 g to separate the soluble proteins from cell debris. A further purification was made by passing the proteins through an illustra NAP-5 column (GE Healthcare Life Sciences, Little Chalfont, Buckinghamshire, UK).

Enzyme activity assays were performed to test the terpene production of the different CsTPS. Activity assays were carried out by adding 50 μL of assay buffer and 50 μL of purified crude bacterial protein extract with 10 mM MgCl_2_ and 10 μM (*E,E*)-FPP into a threaded 1 mL glass vial with a cap containing a Teflon septum. The reaction mix was incubated for 1 h at 30 °C. During the incubation period, VOCs were sampled with a SPME fiber. For volatile analysis, the collected volatiles were desorbed directly in the inlet (240 °C) of the GC-MS system. An Agilent 6890 series GC coupled to an Agilent 5973 MSD (source 230 °C, quadrupole 150 °C, ionization potential 70 eV; Palo Alto, CA, USA) was used for analysis. He was used as carrier gas at a rate of 1 mL*min^−1^. The volatile separation took place on a DB-5MS capillary column (Agilent, Santa Clara, CA, USA, 30 m × 0.25 mm × 0.25 μm). The initial oven temperature of 80 °C was hold for 2 min, followed by a ramp of 7 °C/min to 180 °C and a second ramp of 100 °C/min to 300 °C where the temperature was held for 1 min.

### qRT-PCR analysis of CsTPS genes

To determine the expression levels of individual CsTPS genes, RNA was extracted, converted into cDNA and further used for qRT-PCR. Total RNA was isolated from the same root and leaf tissue samples as for hexane extraction. This was made following the InviTrap^®^ Spin Plant RNA Mini Kit (Stratec molecular, Berlin, Germany). Next, 1 μg of the RNA was DNase I treated followed by first-strand cDNA synthesis using RevertAid H Minus Reverse Trascriptase with oligo (dT)_18_ primers (Thermo scientific, CA, USA). cDNA was diluted 1:10 before used for qRT-PCR. To find an appropriate reverence gene, *actin1* and *EF1α* sequences of *Arabidopsis thaliana* were taken as query for a screen in the *C. stoebe* Trinity assembly with the software Blast2GO 4.1 (Götz et al. 2008) running at default settings. Two primer combinations were designed for each homologous reference gene. *EF1α* was found to be the most robust reverence gene. Next, for each of the CsTPS genes, a qPCR primer pair was designed. All primers are listed in supplementary (Tab. S2). Primer specificity was tested by means of melting curve analysis and gel electrophoresis. qRT-PCR was carried out on a LightCycler^®^ 96 Instrument (Roche, Basel, Switzerland) using the KAPA 480 SYBR FAST qPCR Master Mix (Kapa Biosystems, Boston, USA). Primer efficiency was determined using a linear standard curve approach. For very low expressed genes, this was repeated with samples spiked with plasmids containing the genes of interest. Biological replicates were all run in technical triplicates. Three samples had to be excluded from the analysis due to poor RNA quality or very low expression of the reference gene, resulting in a total of 7 biological replicates for *CsTPS4* as well as *CsTPS5* and 5 biological replicates for *CsTPS1*. Relative transcript abundance was analyzed as fold change (2^-ΔCt^). As *CsTPS1* showed dissimilar melting peaks for root and shoot PCR amplicons, the fragments were subsequently sequenced by Sanger sequencing.

### Impact of C. stoebe root VOCs on neighboring plants

To evaluate the influence of *C. stoebe* root volatiles on the germination and growth of neighboring plants, we used an experimental setup that excluded direct root contact or the transfer of exudates, but allowed *C. stoebe* root VOCs to diffuse to the neighboring plants. The system consisted of mesh cages (12 × 9 × 10 cm, length × width × height) made of Geotex fleece (Windhager, Austria), which were placed in pairs into rectangular plastic pots (Fig. 4A). A covered airgap between the cages allowed for the diffusion of VOCs between the rhizospheres of plants growing in the soil-filled mesh cages. Water was supplied carefully to soil in the mesh cages avoid leaching and exchange of root exudates across the airgap. The Geotex fleece of the mesh cages was sufficient to stop roots from growing out of the mesh cages, thus eliminating direct root contact between the plants. Diffusion of *C. stoebe* VOCs into the airgap was confirmed by SMPE (companion paper Huang et al., under review). Plants for this experiment were grown in a greenhouse (light: 14h; temperature: day 16-24 °C, night 16-22 °C, mean temperature over growth period 20 °C; humidity: 30-60 °C) in potting soil consisting of five parts ‘Landerde’ (RICOTER, Aarberg, Switzerland), four parts ‘Floratorf’ (Floragard, Oldenburg, Germany) and one part sand („Capito“ 1-4 mm, LANDI Schweiz AG, Dotzigen, Switzerland). The “sender” mesh cages in the plastic pots where either left plant free or planted with three week old *C. stoebe* seedlings. After 25 days, different plant species were planted into the “receiver” mesh cages (10 seeds per cage, n = 12 for each species). As receiver species, 11 commonly co-occurring species of *C. stoebe* were selected: *Anthemis tinctoria*, *Cardaria draba*, *Centaurea stoebe*, *Cichorium intybus*, *Dianthus carthusianorum*, *Echium vulgare*, *Festuca valesiaca*, *Koeleria macrantha*, *Medicago sativa*, *Ranunculus bulbosus*, *Daucus carota*, and *Taraxacum officinale* agg. The pots were watered every one to three days. Pots were turned 180° and randomized fortnightly. Potential bias through above ground effects of *C. stoebe* was ruled out by arranging the pots on the table so that each receiver had a *C. stoebe* as neighbor either only above ground in a separate pot (control) or aboveground and belowground in the same pot (treatment). The total number of germinated seeds was recorded after 4 weeks. The first germinated seedling was retained, all the others were removed. After nine weeks of growth, the plants were harvested. Roots and leaves were washed, separated and dried at 80 °C until constant weight to determine dry mass.

### Data Analysis

Statistical assumptions such as normal distribution and homoscedasticity of error variance were checked and square root or log_e_ transformed if the assumptions were not met. Differences in relative peak area per g FW between root and leaf tissue in hexane extracts were tested with a Wilcoxon signed rank test. To test for differences in sesquiterpene abundance among *C. stoebe* populations and *Centaurea* species for a given compound, Analysis of Variance (ANOVA) of a fitted linear model was performed and if significant followed by LS means pairwise comparisons with *p* value adjustment. Differences in expression levels between root and leaf tissue were tested by Wilcoxon signed rank tests. A possible effect of the emitter on the germination was analyzed by fitting a generalized linear model with a quasibinomial distribution to the data and performing an ANOVA (n = 12 per species and treatment). Dry biomass of roots and leaves were investigated by fitting a linear model and conducting an ANOVA (n = 12 per species and treatment, 9 out of 244 plants died and were therefore excluded from the analysis). For each species, the effect of the emitter plant on biomass production was tested by means of a Student’s *t*-test followed by *p* value correction for multiple comparison (Benjamini & Hochberg 1995). Statistical analysis and data visualization was conducted with R 3.4.3 (R Core Team 2017), with ‘lsmeans’, ‘car’ ‘plyer’ and ‘ggplot2’ packages (Lenth 2016; Wickham 2009, 2011; Fox & Weisberg 2011).

## Results

### Characterization of C. stoebe VOCs

Analysis of the volatile blend emitted by intact *C. stoebe* roots revealed an abundant sesquiterpene fraction (Fig. 1A) with (*E*)-β-caryophyllene and daucadiene (most likely a diastereoisomere of trans-dauca-4(11),8-diene) as the predominant compounds. The sesquiterpenes (*E*)-α-bergamotene, humulene, (*E*)-β-farnesene, three putative petasitene isomers (petasitene 1-3), and an unknown sesquiterpene were emitted as well. (*E*)-β-caryophyllene emission was quantified at 3.15 ± 0.69 μg g^−1^ dw h^−1^ (mean ± SE). Hexane root tissue extracts contained comparable sesquiterpene profiles, with (*E*)-β-caryophyllene and daucadiene as major compounds (Fig. 1B). Additionally, low quantities of other sesquiterpenes such as cyclosativene, β-acoradiene, α-farnesene, and β-bisabolene were found in these extracts, which were not detected in the volatile blend of intact roots. Besides sesquiterpenes, there were other compounds eluting from the column, mostly at later time points. The most abundant of these compounds showed a terpenoid-like structure and was tentatively identified as a sesquiterpene lactone (*m/z* = 232). The other late eluting analytes were neither known nor present in the volatile blend of intact roots and therefore not analyzed further. Sesquiterpenes were much more abundant in root hexane extracts than leaf extracts (Fig. 1B). Only four compounds were detected in both leaves and roots, namely α-copaene, (*E*)-β-caryophyllene, δ-cadinene and the putative sesquiterpene lactone. (*E*)-β-caryophyllene and the putative sesquiterpene lactone were present in much higher concentrations in the roots than the leaves (Wilcoxon signed rank test: n = 10, *p* = 0.002), while α-copaene, δ-cadinene were more abundant in the leaves (Wilcoxon signed rank test: n = 10, *p* = 0.002). In contrast to root tissue, we also detected three monoterpenes in *C. stoebe* leaves: α-pinene, β-myrcene and an unknown monoterpene. Compared to sesquiterpenes, monoterpene signals were low in abundance.

**Fig. 1.**
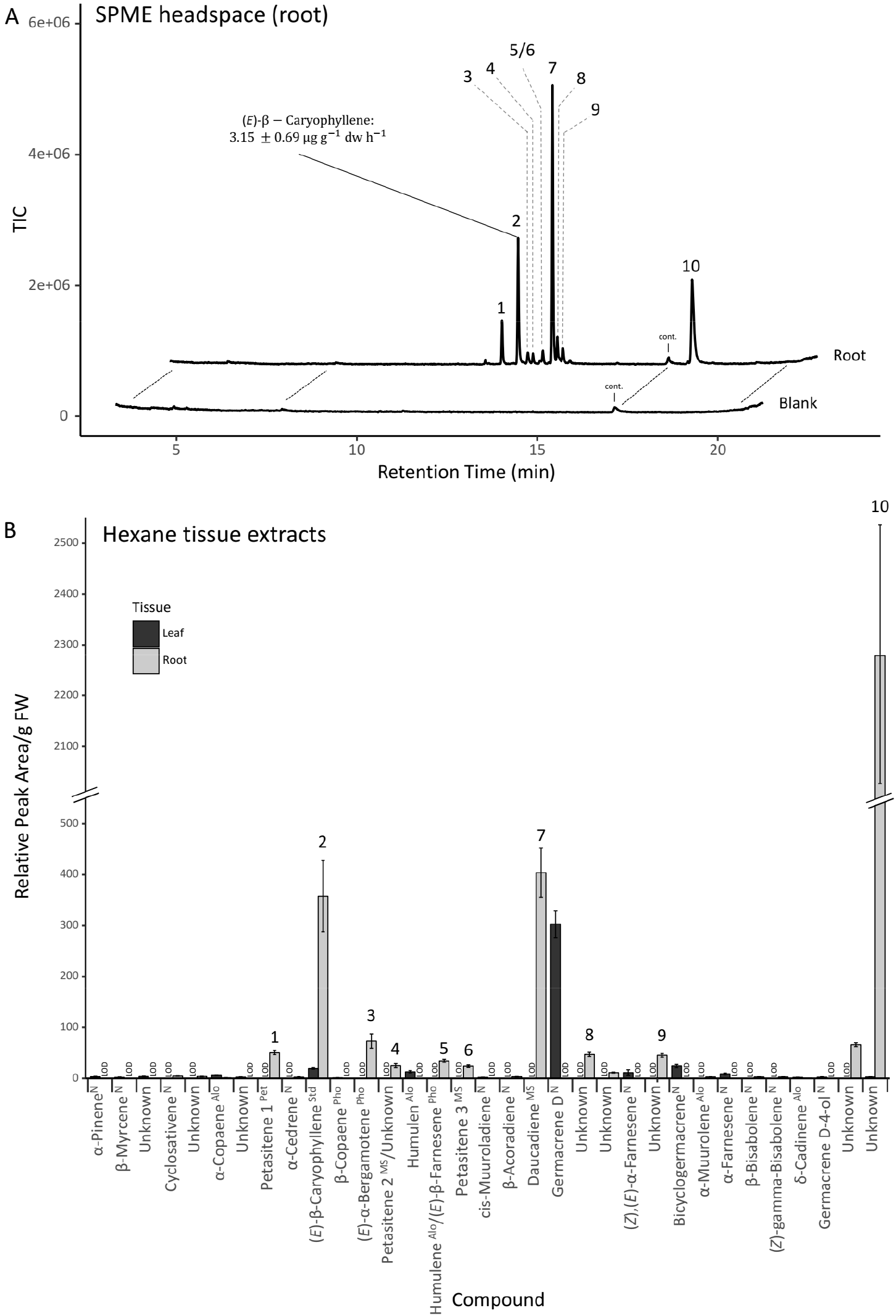
Centaurea stoebe roots release high amounts of sesquiterpenes. (A) Representative SPME-GC-MS chromatogram of VOCs emitted by intact *C. stoebe* roots. (*E*)-β-caryophyllene emission rate is displayed as mean ± SE (n = 8; dw, dry weight). (B) Relative peak area per g fresh weight (FW) of compounds found in hexane tissue extracts shown as mean ± SE (n = 10). TIC, total ion current; 1, petasitene 1; 2, (*E*)-β-caryophyllene; 3, (*E*)-α-bergamotene; 4, petasitene 2; 5, humulene and (*E*)-β-farnesene; 6, petasitene 3; 7, daucadiene; 8, unknown sesquiterpene; 9, unknown non terpenoid; 10, unknown sesquiterpene lactone-like compound; cont, contamination; LOD, below limit of detection; Identification: N, NIST library, comparison of mass spectra and retention index (RI); MS, inspection of mass spectra (RI other than literature); Std, comparison of mass spectra an RI with pure standard compound; and comparison of mass spectra an RI with known compounds of Alo, *Aloysia sellowii*; Pet, *Petasites hybridus;* Pho, *Phoebe porosa*.

### Emission Pattern of C. stoebe Populations and other Centaurea Species

Sesquiterpenes released by intact roots of three different *C. stoebe* populations did not differ significantly in quality and quantity (Fig. 2A), suggesting that this trait is conserved within *C. stoebe*. By contrast, congeneric *Centaurea* species emitted distinct terpene bouquets compared to *C. stoebe* (Fig. 2B). The volatile blend of the closely related *C. valesiaca* was most similar to *C. stoebe*, with petasitene 1, petasitene 2 and daucadiene being emitted in lower quantities by *C. valesiaca* than by *C. stoebe. C. jacea* emitted sesquiterpenes similar to *C. stoebe* but in different quantities: the release of petasitene 1, petasitene 3, (*E*)-α-bergamotene and of an unknown compound was significantly increased in *C. jacea* compared to *C. stoebe*. Finally, we detected (*E*)-β-caryophyllene and (*E*)-α-bergamotene, but not any of the other sesquiterpenes in the headspace of *C. scabiosa* roots. Thus, sesquiterpene release from the roots seems to be conserved in *C. stoebe* ecotypes, but varies qualitatively and quantitatively between different *Centaurea* species.

**Fig. 2.**
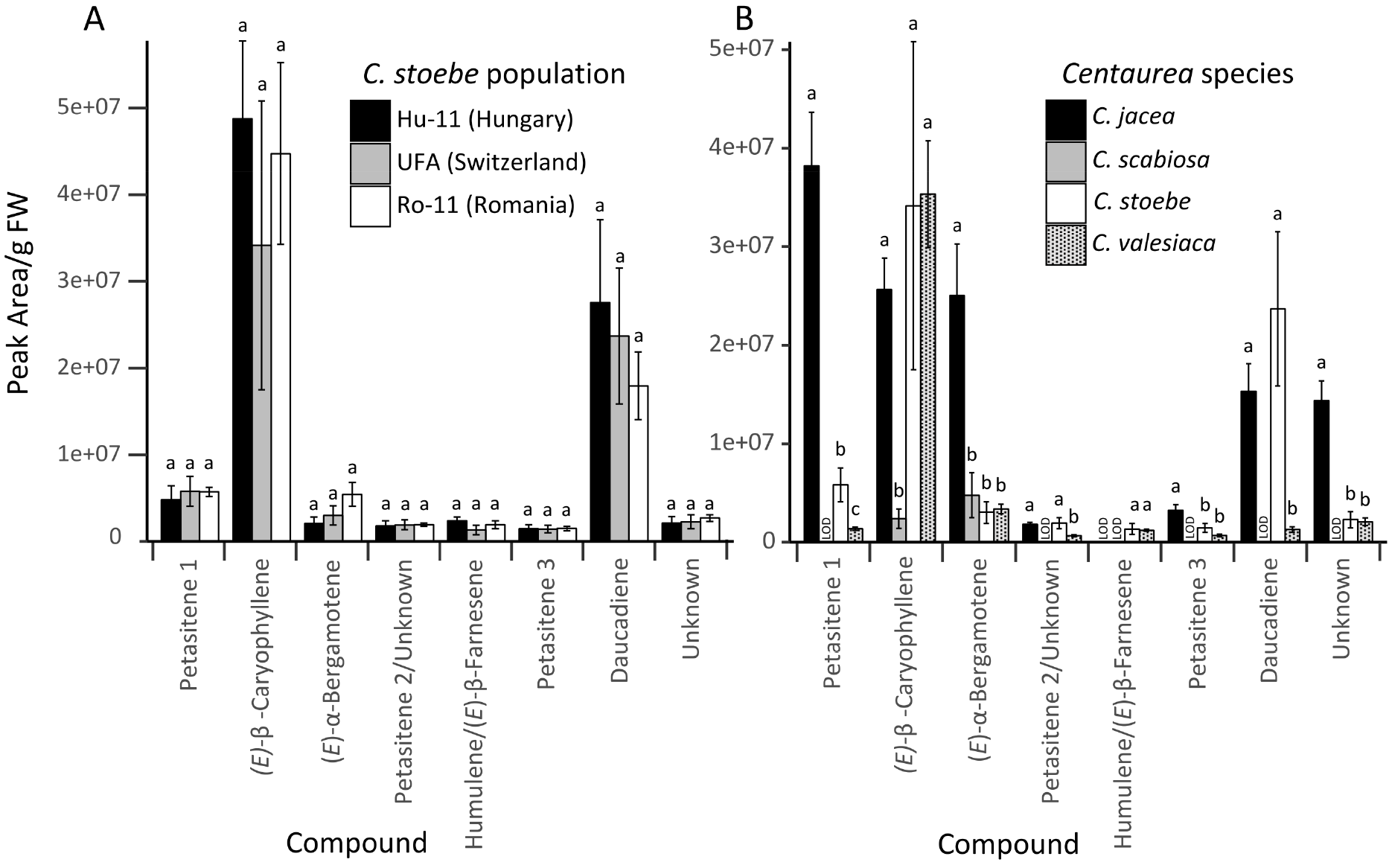
Root sesquiterpene release is conserved within *C. stoebe*, but varies between different *Centaurea* species. (A) Peak area per g fresh weight (FW) of *C. stoebe* populations shown as mean ± SE (n = 5; except for Hu-11, n = 7). Letters show significant differences among populations within one compound (Analysis of Variance (ANOVA) followed by pairwise comparison of LS means, *p_adj_* < 0.05). (B) Peak area per g fresh weight (FW) of *Centaurea* species shown as mean ± SE (*C. jacea* and *C. scabiosa*, n = 8; *C. stoebe*, n = 5; *C. valesiaca*, n = 4). Letters show significant differences among species within one compound (Analysis of Variance (ANOVA) followed by pairwise comparison of LS means, *p_adj_* < 0.05). LOD, below limit of detection.

### Terpene Synthases of C. stoebe

To understand the genetic basis of sesquiterpene formation in *C. stoebe* roots, known sequences of *M. recutita* terpene synthases (TPS) were used to find homologous genes in the *C. stoebe* root transcriptome. This led to the identification of eight potential sesquiterpene synthases (CsTPSs, Fig. 3A). Apart from CsTPS2 and CsTPS3, for which ORF amplification and transformation into *E. coli* was unsuccessful, all TPSs were successfully cloned and expressed in *E. coli*. CsTPS protein activity assays showed that CsTPS1, CsTPS4, CsTPS5, CsTPS7, and CsTPS8 exhibit sesquiterpene synthase activity. No activity was found for CsTPS6 (Fig. 3B-G). CsTPS1 catalyzed the formation of α-muurolene, and CsTPS4 produced (*E*)-β-caryophyllene and humulene. CsTPS5 produced daucadiene as main compound and (*E*)-α-bergamotene, (*E*)-β-farnesene, three petasitenes, β-acoradiene, β-bisabolene, (*Z*)-γ-bisabolene, as well as an unknown sesquiterpene as byproducts. All the compounds produced by CsTPS1, CsTPS4 and CsTPS5 were found in hexane root extracts of *C. stoebe*. Furthermore, the compounds produced by CsTPS4 and CsTPS5 cover all highly emitted volatiles from intact roots. Comparison of retention indices and mass spectra revealed that CsTPS7 produced (*E*)-α-bisabolene (RI 1545) and CsTPS8 produced α-zingiberene (RI 1497) as main compounds. The two compounds were not detected in tissue extracts or the headspace of intact roots.

**Fig. 3.**
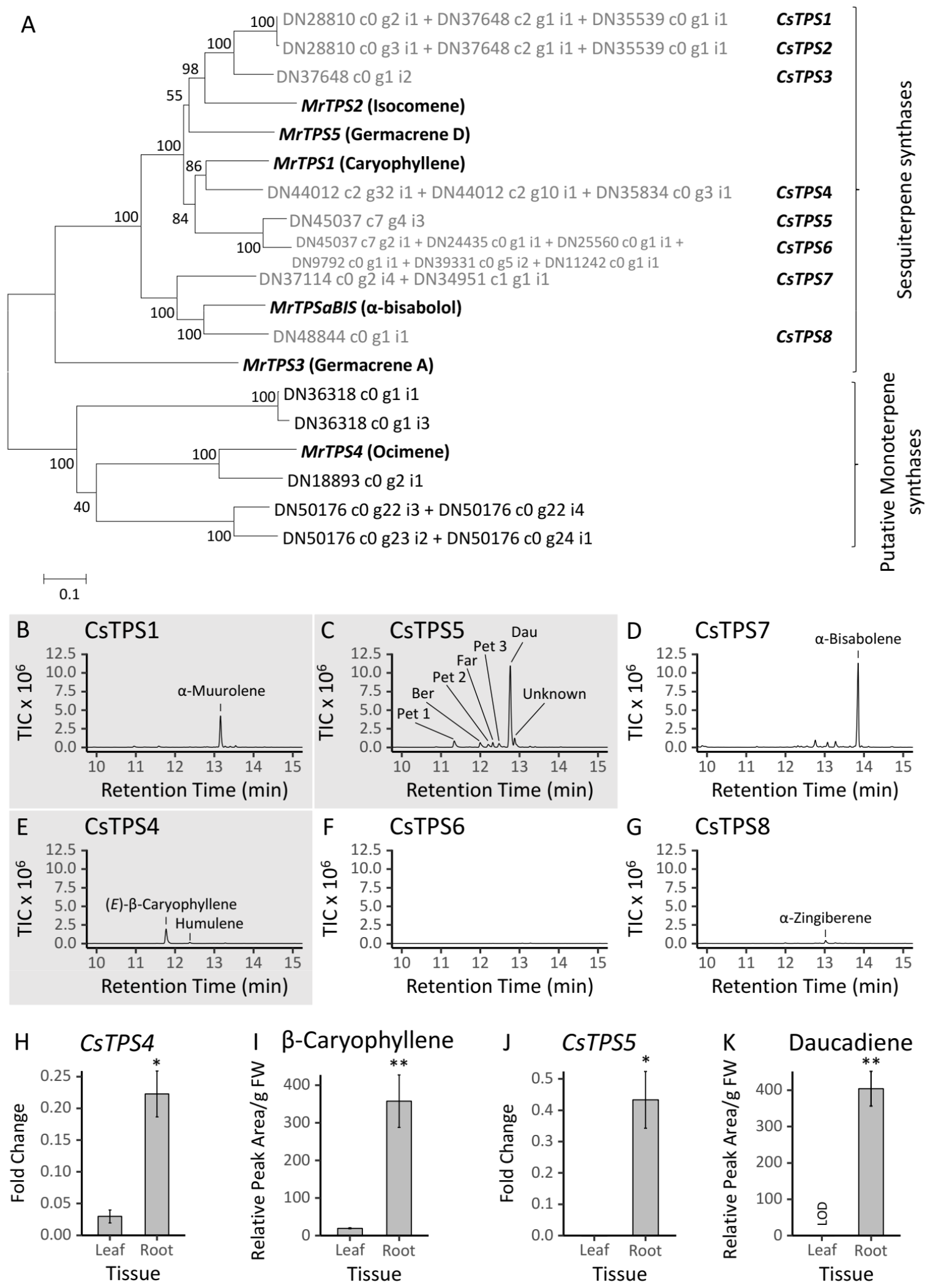
Two terpene synthases account for major *Centaurea stoebe* root sesquiterpenes. (A) To find potential *C. stoebe* terpene synthases (CsTPSs), sequences of *Matricaria recutita* terpene synthases (MrTPS) were taken to screen for homologous genes in the *C. stoebe* root transcriptome. The phylogenetic tree shows contigs of potential *CsTPSs* as end nodes and their related MrTPS genes. (B-G) SPME-GC-MS analysis of CsTPS protein activity assays with (*E,E*)-FPP as substrate. Compounds of highlighted chromatograms (B, C, E) were also found in *C. stoebe* hexane root extracts. mRNA abundance for *CsTPS4* (H) and *CsTPS5* (J) and relative peak area per g fresh weight (FW) of their main products (*E*)-β-caryophyllene (I) and daucadiene (K) in hexane root extracts. Shown are mean ± SE (qRT-PCR, n = 7; Tissue extracts, n = 10). Differences in means were tested by Wilcoxon signed rank tests, levels of significance: *p* < 0.01 **, *p* < 0.05 *. TIC, total ion current; Pet, petasitene; Ber, (*E*)-α-bergamotene; Far, (*E*)-β-farnesene; Dau, daucadiene. LOD, below limit of detection

The predominant sesquiterpenes (*E*)-β-caryophyllene and daucadiene are produced in high amounts in the roots, but not in the leaves (Fig. 3I/K). The same pattern was found for the expression of *CsTPS4* and *CsTPS5*, the two TPSs putatively responsible for the production of these VOCs (Fig. 3H/J). The mRNA levels in root compared to leave tissue revealed a 7.5-fold increase in *CsTPS4* (Wilcoxon signed rank test: n = 7, *p* = 0.016) and a >5,000-fold increase for *CsTPS5* (Wilcoxon signed rank test: n = 7, *p* = 0.016). Low expression of *CsTPS1* was detected in the leaves and roots. Melting point analysis indicated that different fragments were amplified in the different tissues. Fragment sequencing revealed that the root fragment corresponds to *CsTPS1*, whereas the leaf fragment only showed 89% sequence similarity to *CsTPS1*. No other sequence in the *C. stoebe* root transcriptome besides *CsTPS1* was found to match the leaf fragment, suggesting that it may stem from a TPS gene that is specifically expressed in the leaves.

### Effect of C. stoebe Root Volatiles on Neighboring Plants

To test whether *C. stoebe* root VOCs influence the germination and performance of neighboring plants, we exposed seeds and germinating plants of different sympatric species to *C. stoebe* rhizosphere VOCs for several weeks. An overall positive effect of *C. stoebe* root VOCs on the germination of the different sympatric plant species was observed (*p* = 0.03, Fig. 4B). Furthermore, nine weeks after sowing, root biomass (*p* = 0.02, Fig. 4C) and leaf biomass (*p* = 0.006, Fig. 4D) were significantly increased in the presence of *C. stoebe* root VOCs. Individual comparisons revealed no significant effects for single species, even though visual inspection indicated that the magnitude of the effects varied from neutral to positive for the different species.

**Fig. 4.**
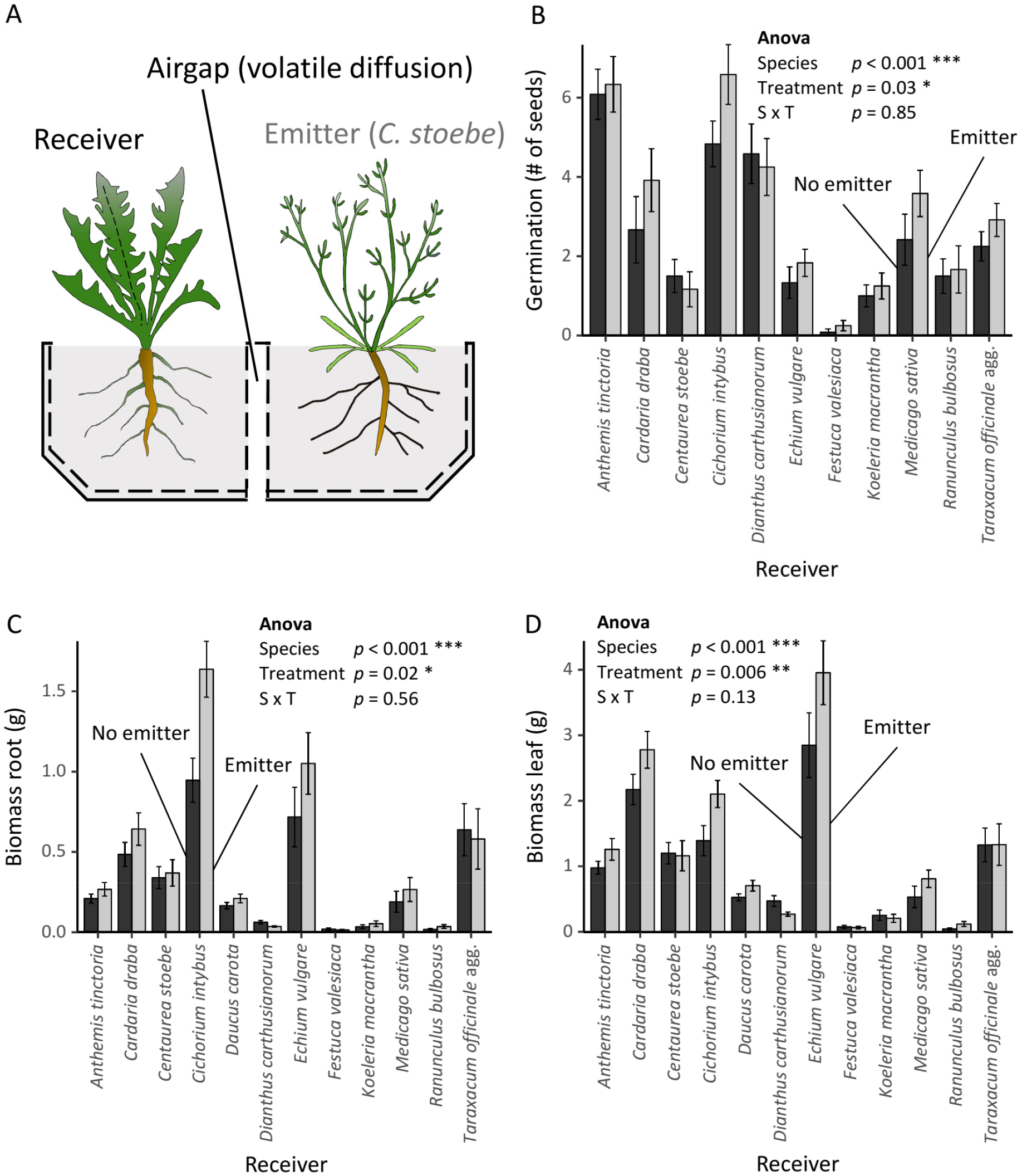
*Centaurea stoebe* root volatiles increase germination and growth of sympatric neighbors. (A) Experimental setup to evaluate the influence of *C. stoebe* (‘emitter’) root volatiles on receiver plant species. As control, the emitter compartment was filled with soil, but no plant was grown in it (‘no emitter’). (B) Number of receiver seeds that germinated up to four weeks after they were sown. Analysis of Variance (ANOVA) output of generalized linear model is shown (distribution, quasibinomial; n = 11 per species and treatment). Dry biomass of receiver roots (C) and leaves (D) after nine weeks of growth. ANOVA output of linear model is shown (levels of significance: *p* < 0.001 ***, *p* < 0.01 **, *p* < 0.05 *; n = 12 per species and treatment)

## Discussion

Plants are known to produce a variety of VOCs that play important roles in biotic interactions (Peñuelas et al. 2014; Pichersky et al. 2002). Physiological changes in plants exposed to VOCs from neighboring plants for instance are well documented above ground (Arimura, Shiojiri & Karban 2010; Heil & Karban 2010; Karban et al. 2014). In contrast, there is a gap of knowledge regarding VOC-mediated plant-plant interactions below ground (Delory et al. 2016). In this study, we characterized the volatiles emitted by *C. stoebe* and identified two terpene synthases which are sufficient to produce the full sesquiterpene blend emitted by intact roots. Furthermore, we show that *C. stoebe* root VOCs enhance germination and biomass production of sympatric neighbors. Here, we discuss these findings from physiological and ecological points of view and reflect on the potential role of root VOCs in determining the rarity of *C. stoebe* in its native environment.

Plants can release terpenoids constitutively or in response to environmental stress (Keeling & Bohlmann 2006). Our headspace analyses show that *C. stoebe* releases sesquiterpenes specifically and constitutively from its roots. The emission rate of the sesquiterpene (*E*)-β-caryophyllene was measured at 3.15 ± 0.69 μg g^−1^ dw h^−1^ (mean ± SE), leading to a situation where 2 seconds of exposure to a few mg of *C. stoebe* roots already saturated our analytical equipment. For comparison, (*E*)-β-caryophyllene release from herbivore-attacked maize roots is likely in the lower ng range per plant (Hiltpold et al. 2011). Only few studies so far provide absolute quantification of root VOC emission rates, and we are not aware of any report showing below ground sesquiterpene release rates at the levels reported here. Monoterpenes have been shown to be released in substantial quantities by roots. *Pinus pinea* roots for instance release monoterpenes at rates up to 26 ± 5 μg g^−1^ dw h^−1^ (mean ± SE) (Lin, Owen & Peñuelas 2007). Thus, *C. stoebe* constitutively releases relatively high amounts of sesquiterpenes from its roots.

Terpenoids are produced by terpene synthases (TPSs) (Bohlmann, Meyer-Gauen & Croteau 1998). We identified two CsTPSs whose products correspond to the root-emitted sesquiterpenes in *C. stoebe*. (*E*)-β-caryophyllene occurs in many plant species and it has been reported several times to be produced by the same terpene synthase as humulene (Cai et al. 2002; Irmisch et al. 2012; Köllner et al. 2008; Yang et al. 2013). In *C. stoebe*, we also found these two compounds to be produced by the same TPS (CsTPS4). Examining the expression level of *CsTPS4* in roots and leaves of *C. stoebe* showed the same pattern as the distribution of the compound: low quantities of RNA and (*E*)-β-caryophyllene in leaves and significantly higher quantities of both in roots. The second TPS involved in producing the volatile bouquet is CsTPS5 with daucadiene as main product. Enzyme activity assays of this enzyme led to the production of several sesquiterpenes, all of which were also present in *C. stoebe* roots. The sesquiterpenes produced by CsTPS5 were not found in the leaves, and *CsTPS5* was not expressed in this tissue. Regulation of sesquiterpene synthesis through transcriptional control of *TPSs* is well established (Tholl 2006) and likely also accounts for the differences in leaf and root sesquiterpene profiles in *C. stoebe*. Taken together, we show that two highly expressed, root-specific TPSs can account for the full root sesquiterpene blend of *C. stoebe*.

*In vitro* studies found negative effects of root VOCs on seed germination (Ens et al. 2009; Jassbi et al. 2010). Using a soil-based system that allows for the passive diffusion of VOCs between sender and receiver plants, we demonstrate that *C. stoebe* volatiles have no negative effects on the germination and growth of 11 sympatric plant species. Root VOC exposure even resulted in an overall increase in the germination and growth of other plants. A degradation product of (*E*)-β-caryophyllene has been shown to exhibit a broad antifungal activity (Hubbell, Wiemer & Adejare 1983) and other root VOCs are also known to influence microbial communities, which again can alter plant performance (Wenke et al. 2010; Inderjit & Weiner 2001; Kleinheinz et al. 1999). Thus, the positive effect of *C. stoebe* root VOCs on the receiver plants could either be a direct effect mediated through the impact of the VOCs on the physiology of the seeds and growing plants, or an indirect effect mediated through soil microbial communities (Hu et al. 2018b). Of note, *C. stoebe* VOCs do not only modulate plant performance, but can also change root physiology and herbivore resistance, as shown in the companion paper to this study (companion paper Huang et al., under review). Thus, the effects of *C. stoebe* VOCs on neighboring plants are likely multifaceted and may change the interactions of neighboring plants with other organisms. How root VOCs interact with bioactive soluble exudates, which can also be important for plant and herbivore performance (Hu et al. 2018a), remains to be studied.

The release of VOCs can benefit the emitter by intoxicating and repelling herbivores, attracting natural enemies and priming defenses in systemic tissues (De Moraes, Mescher & Tumlinson 2001; Frost, Mescher, Carlson, De Moraes 2008; Erb et al. 2015; Schuman, Barthel & Baldwin 2012; Ye et al. 2018). To what extent the release of VOCs is beneficial for the emitter in the context of plant-plant interactions, however, is less clear. Here, we show that the release of sesquiterpenes from the roots may have negative consequences for *C. stoebe* plants, as it increases the germination and growth of a variety of sympatric competitors. Strikingly, and in contrast to what has been observed in other plant systems (Degen et al. 2004; Schuman et al. 2009), sesquiterpene release seems to be conserved within different *C. stoebe* ecotypes. The benefit of this potentially conserved phenotype for *C. stoebe* is currently unclear. Germination and growth of *C. stoebe* itself does not seem to be improved through VOC exposure, for instance. However, it is possible that the high release rates protect the plant from herbivores and pathogens in addition to the known resistance factors in this species (Landau, Müller-Schärer & Ward 1994). Furthermore, as shown in the companion paper (companion paper Huang et al., under review), the VOCs may trigger susceptibility to herbivores in neighboring species. Knocking down *CsTPS4* and *CsTPS5* could help to understand the potential benefits of root sesquiterpene production in the future.

According to the IUCN red list, *C. stoebe* is classified as threatened in Switzerland while it is invasive in the United States. Substantial work has been conducted to evaluate whether *C. stoebe* may suppress competitors in the invasive range through allelopathic effects (Duke et al. 2009; Ridenour & Callaway 2001). It has for instance been demonstrated that *C. stoebe* suffers substantially from competition by its neighbors in its native range, but not in the invasive range (Callaway et al. 2011). It will be interesting to study VOC emissions of invasive ecotypes and effects on competitors in the invasive range in the future. In the native range, the increased growth of neighboring species triggered by *C. stoebe* root VOCs may contribute to its rarity.

In conclusion, this work demonstrates that two TPSs are sufficient to explain the high constitutive sesquiterpene emissions of *C. stoebe*, and that the release of these VOCs, as dominant constituents of the full root VOC blend, do not negatively affect neighboring plants, but increase their growth and germination. Thus, below ground plant-plant interactions mediated by plant volatiles may affect competition and coexistence in natural plant communities.

## Acknowledgements

We thank Adrian Möhl (Info Flora) for advice on plant species growing in sympatry with *C. stoebe*. Additionally, we thank Yan Sun and Heinz Müller Schärer (University of Fribourg) as well as Adrian Möhl and Markus Fischer (University of Bern) for providing seeds. This project was supported by the European Commission (MC-IEF no. 704334 to W.H.) and the University of Bern.

## Authors Contributions

V.G, T.G.K. and M.E. designed the experiments. M.H. and T.G.K. sequenced, assembled and analyzed the *C. stoebe* root transcriptome. V.G. and C.F. performed experiments. V.G., T.G.K. and M.E. analyzed data. V.G. and M.E. wrote the first draft of this manuscript.

## Data accessibility

Raw data associated with this study can be downloaded from Dryad [to be inserted at a later stage] and the NCBI Sequence Read Archive (SRA) [to be inserted at a later stage].

## Supplementary materials

**Table S1:**
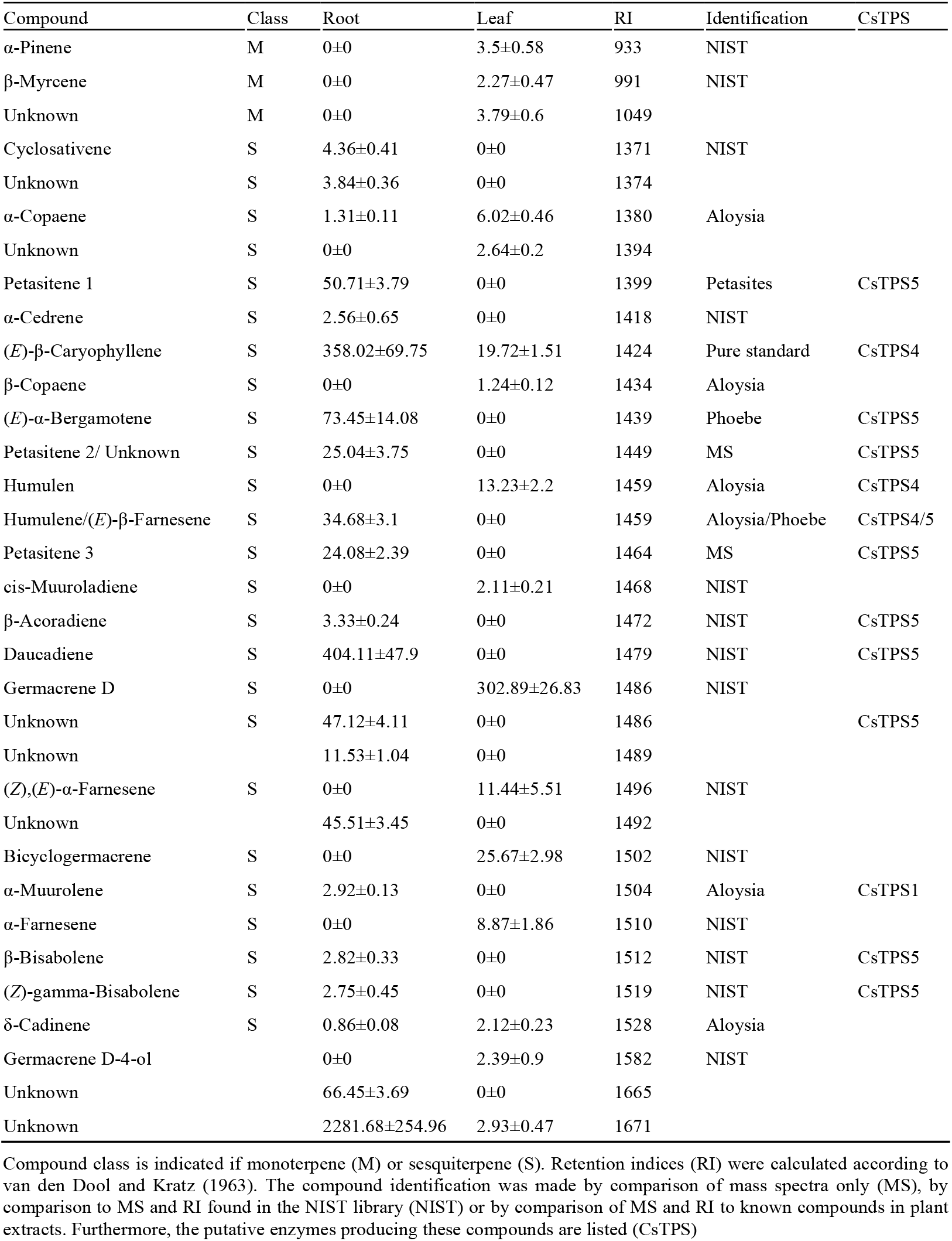
Compounds found in *Centaurea stoebe* hexane tissue extracts listed as means ± SE of relative peak area per g fresh weight.

**Table S2:**
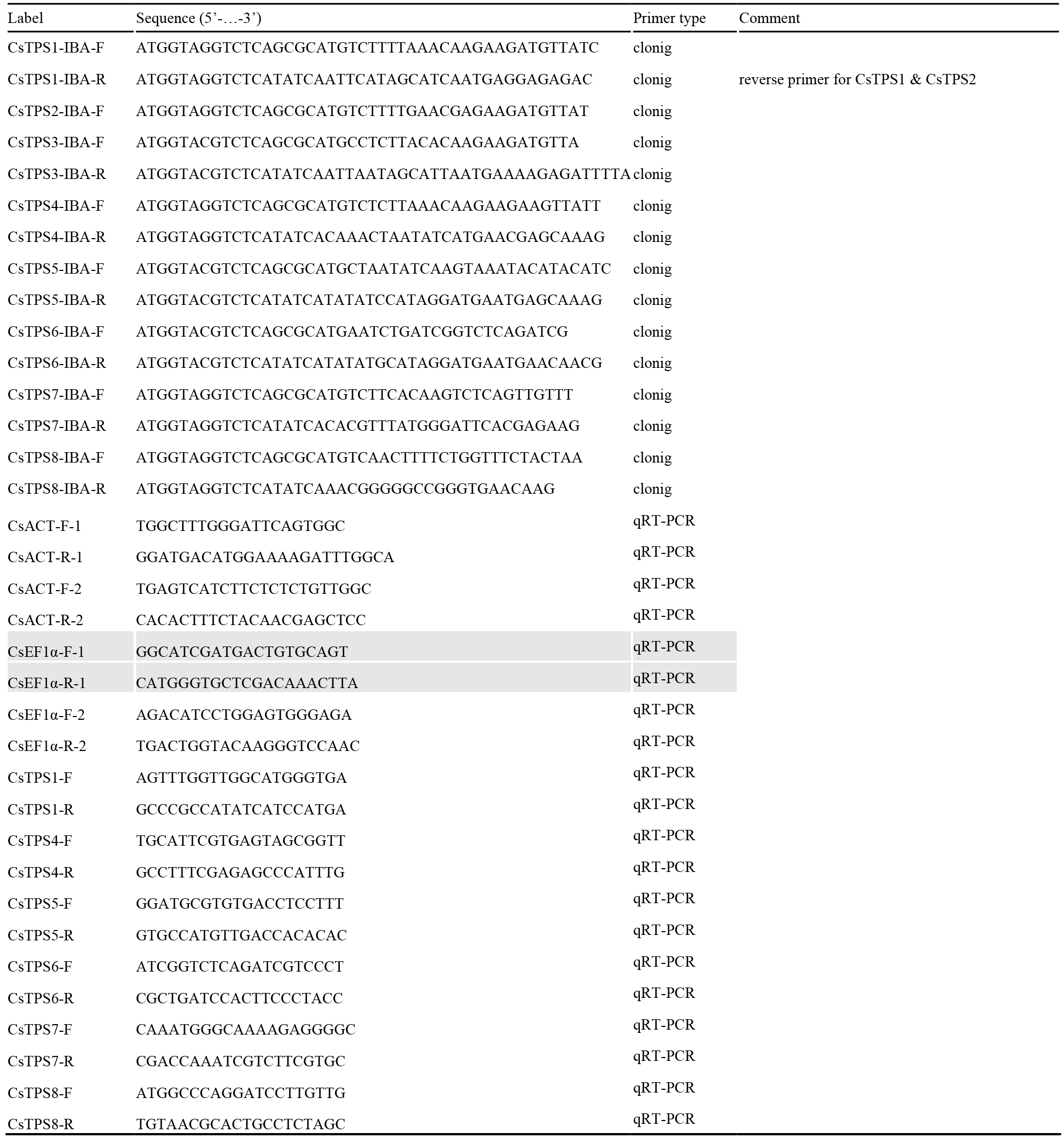
Primers used for cloning of *CsTPS* genes and for qRT-PCR. The reverence gene primer pair used for the analysis is highlighted in grey.

